# Tidal volume and respiration phase modulate cortico-muscular communication

**DOI:** 10.1101/2020.01.13.904524

**Authors:** Daniel S. Kluger, Joachim Gross

## Abstract

Recent studies in animals have convincingly demonstrated that respiration cyclically modulates oscillatory neural activity across diverse brain areas. To what extent this generalises to humans in a way that is relevant for behaviour is yet unclear. We used magnetoencephalography (MEG) to assess the potential influence of tidal volume and respiration phase on the human motor system. We obtained simultaneous recordings of brain activity, muscle activity, and respiration while participants performed an isometric contraction task. We used corticomuscular coherence as a measure of efficient long-range cortico-peripheral communication.

We found coherence within the beta range over sensorimotor cortex to be reduced during voluntary deep compared to involuntary normal breathing. Moreover, beta coherence was found to be cyclically modulated by respiration phase in both conditions. Overall, these results demonstrate how respiratory rhythms actively influence brain oscillations in an effort to synchronise neural activity for the sake of computational efficiency. Intriguing questions remain with regard to the shape of these modulatory processes and how they influence perception, cognition, and behaviour.

## Introduction

The act of breathing is one of the fundamental and vital rhythms of life. In humans at rest it occurs at a frequency of about 0.25 Hz and consists of active inspiration and passive expiration (Fleming et al., 2011). Interestingly, breathing is temporally coordinated by coupled oscillators that provide the periodic drive to initiate inspiration. Central among these generators of the breathing rhythm is a microcircuit in the medulla, the preBötzinger complex. It typically controls breathing autonomously. However, respiration is also under top-down control. In fact, it plays an essential role in a range of human behaviours such as speaking, singing, laughing and crying (McKay et al., 2003). The relationship between respiration and higher cognitive functions, however, goes far beyond temporal coordination of the breathing act. Cortical brain areas receive afferent respiratory input from the preBötzinger complex (e.g. via the locus coeruleus), from olfactory nuclei and through the vagus nerve (Del Negro et al., 2018). In turn, brain states (such as anxiety) lead to changes in respiration.

The mechanisms underlying these bi-directional interactions and their effects on cognition are not fully understood, whereas it has been reported that a range of cognitive and motor functions change with the phase within the respiratory cycle. For example, participants spontaneously inhale at onsets of cognitive tasks and visuospatial behavioural performance is better during inspiration compared to expiration (Perl et al., 2019). Respiration also cyclically modulates the magnitude of the acoustic startle response (Schulz et al., 2016), the amplitude of regionally specific brain activity and memory performance for fear stimuli (Zelano et al., 2016). In addition, maximum muscle force (Li & Laskin, 2006), oculomotor control (Rassler & Raabe, 2003), near-threshold perception (Flexman et al., 1974), and face processing (Zelano et al., 2016) depend on the phase within a respiratory cycle. This bi-directional interaction between respiration and brain processing is markedly characterised by the joint feature of rhythmicity: Brain rhythms as sensitive markers of brain states have widely been described across cognitive domains including attention, perception, and memory (Thut et al., 2012; Wang 2010). However, it remains unclear how rhythmic respiration and rhythmic brain activity mutually interact.

The fundamental importance of this research question is highlighted by recent evidence from the animal literature demonstrating that breathing entrains brain oscillations not only in olfactory regions (Rojas-Líbano et al., 2014; Frederick et al., 2016) but also in non-olfactory areas like whisker barrel cortex (Moore et al., 2013; Ito et al., 2014) and the hippocampus (Yanovsky et al., 2014; Nguyen Chi et al., 2016). In other words, brain rhythms previously attributed to cognitive processes such as memory were shown to at least in part reflect respiration-related processes (Tort et al., 2018). However seminal, these findings should come to little surprise given well-documented links between respiration and changes in animal behaviour, such as exploration (Welker, 1964; Verhagen et al., 2007) and whisking (Cao et al., 2012) in rodents as well as wing beats and echolocation in bats (Wong & Waters, 2001; Suthers et al., 1972).

Thus, while there is a solid body of evidence illustrating respiration-related modulation of brain oscillations in animal studies, this topic is virtually unstudied in humans. Two recent studies using invasive recordings in epilepsy patients have demonstrated that brain oscillations at various frequencies can be locked to the respiratory cycle even in non-olfactory brain areas (Herrero et al., 2018; Zelano et al., 2016). Moreover, the role of synchronised oscillatory activity was highlighted in an iEEG study conclusively demonstrating how coherence between olfactory and auditory cortices benefits multisensory integration (Zhou et al., 2019). Finally, two of the very few non-invasive accounts to date recently linked respiration phase to changes in task-related oscillatory activity (Hsu et al., 2019) and behavioural measures in a visuospatial task (Perl et al., 2019).

Its functional relevance in multisensory integration notwithstanding, the role of oscillatory coherence (or synchrony) is by no means restricted to the cognitive domain. In fact, coherent oscillations have been suggested to universally mediate and enhance neural communication across distributed neural networks required for current task demands (Engel et al., 2001; Fries, 2005; Fries, 2015). Specifically, oscillations of different frequency ranges are thought to enable dynamic interactions of brain areas at varying spatial scales: While fast oscillations such as gamma rhythms allow functional coupling of neural populations over short distances (e.g., neighbouring cortical regions), slower oscillations like beta rhythms support such neural communication over longer distances (e.g., sensorimotor processing; Kopell et al., 2000). In the motor domain, corticomuscular coherence is a well-established, robust measure of functional connectivity between motor cortex and contralateral effector muscles, characterised by phase synchrony of the two signals (Baker et al., 1999; Brown, 2000; Murthy & Fetz, 1992). This synchrony is best observed during tasks involving isometric contraction, i.e. steady-state motor output (Schnitzler et al., 2000; Feige et al., 2000; Gross et al., 2000; Bourguignon et al., 2019). Corticomuscular coherence (CMC), particularly in the beta band (10 - 30 Hz), has commonly been interpreted as a marker of efficient coding in long-range, bi-directional interaction of the brain and effector muscles (Kristeva et al., 2007; Baker, 2007).

Consolidating these accounts with the initial questions regarding the influence of breathing on neural oscillations, we assessed potential modulatory effects of respiration on CMC as a measure of long-range neural communication. We acquired MEG data from human subjects who lifted and held their right forearm while they executed either automatic (i.e., normal) or voluntary deep breathing. While we hypothesised significant beta-range coherence of muscle and contralateral sensorimotor activity for both deep and normal breathing, we assumed coherence amplitude to differ significantly between the respiratory conditions. Specifically, voluntary deep breathing is associated with active, top-down control of respiratory muscles compared to the more passive, involuntary normal breathing. The active, cortical control of respiration is expected to lead to another corticomuscular communication channel independent of the corticomuscular communication channel subserving isometric arm muscle contraction. In MEG data this is expected to lead to reduced coherence between motor cortex and muscle activity due to the interference from another independent channel. Furthermore, based on the above-mentioned reports of respiration phase modulating a variety of perceptual and cognitive processing, we hypothesised CMC to not only vary as a function of tidal volume (i.e., deep vs normal breathing) but also to be modulated by the phase of the respiration cycle.

## Methods

### Participants

Twenty-eight volunteers (14 female, age 24.8 ± 2.87 years [mean ± SD]) participated in the study. All participants denied having any respiratory or neurological disease and gave written informed consent prior to all experimental procedures. The study was approved by the local ethics committee of the University of Münster and complied with the Declaration of Helsinki.

### Procedure

Participants were seated upright in a magnetically shielded room while we simultaneously recorded MEG, respiration, and Electromyography (EMG) data. MEG data were acquired using a 275 channel whole-head system (OMEGA 275, VSM Medtech Ltd., Vancouver, Canada) at a sampling frequency of 600 Hz. During recording, participants were instructed to keep their eyes on a fixation cross centred on a projector screen placed in front of them. To minimise head movement, participants’ heads were stabilised with cotton pads inside the MEG helmet. Data were acquired in runs of 5 min duration with intermediate self-paced breaks. Alternating between runs, participants were instructed to either breathe normally or voluntarily deeply while maintaining their normal respiration frequency. Tidal volume was measured as thoracic circumference by means of a respiration belt transducer (BIOPAC Systems, Inc., Goleta, United States) placed around the participants’ chest. We recorded two runs per respiratory condition (*normal, deep*), resulting in a total of 4 runs. In each run, participants performed isometric contraction by raising their right forearm by about 30° and stretching all fingers of their open hand slightly upward while resting their elbow on a padded armrest. Corresponding muscle activity was recorded from the posterior right forearm by positioning EMG electrodes over the superficial extensor (*m. extensor carpi radialis longus*) and a reference location on the wrist, respectively. Between runs, participants laid down their right forearm on a padded armrest.

For MEG source localisation we obtained high-resolution structural MRI scans in a 3T Magnetom Prisma scanner (Siemens, Erlangen, Germany). Anatomical images were acquired using a standard Siemens 3D T1-weighted whole brain MPRAGE imaging sequence (1 x 1 x 1 mm voxel size, TR = 2130 ms, TE = 3.51 ms, 256 x 256 mm field of view, 192 sagittal slices). MRI measurement was conducted in supine position to reduce head movements and gadolinium markers were placed at the nasion as well as left and right distal outer ear canal positions for landmark-based co-registration of MEG and MRI coordinate systems.

### MEG and EMG data preprocessing

Data preprocessing was performed using Fieldtrip (Oostenveld et al., 2011) running in Matlab R2018b (The Mathworks, Inc., Natick, United States). We first employed independent component analysis (ICA) to detect and reject cardiac artefacts from the MEG time course. In a second step, we applied a discrete Fourier transform (DFT) filter to eliminate 50 Hz line noise from the continuous MEG data. Finally, EMG data were highpass-filtered at 10 Hz.

### MRI co-registration

Co-registration of structural MRIs to the MEG coordinate system was done individually by initial identification of three anatomical landmarks (nasion, left and right pre-auricular points) in the participant’s MRI. Using the implemented segmentation algorithms in Fieldtrip and SPM12, individual head models were constructed from anatomical MRIs. A solution of the forward model was computed using the realistically shaped single-shell volume conductor model (Nolte 2003) with a 5 mm grid defined in the MNI template brain (Montreal Neurological Institute, Montreal, Canada) after linear transformation to the individual MRI.

### CMC in continuous data

In order to identify MEG sensors of interest (ROI) to be used in our main analyses, we first computed CMC on continuous MEG data. To this end, both runs of either respiratory condition (*normal, deep*) were concatenated for each participant and subsequently segmented into short data segments (duration 1 s, overlap 0.5 s). Spectral power and cross-spectral density of these data were calculated using a single Hanning taper (range 1 - 60 Hz). Coherence between the EMG signal and all MEG channels within the beta band (10 - 30 Hz) was then calculated for both respiratory conditions and averaged across participants.

### CMC in respiration-locked data

To assess potential modulatory effects of respiration phase on cortico-muscular communication, our main analyses focussed on coherence of EMG and MEG signals phase-locked to the respiration cycle. Therefore, we first identified peaks within the respiration signal and calculated the distribution of peak-to-peak distances (i.e., full-cycle duration λ) per subject, respiratory condition, and experimental run. We next used the mean duration from that distribution to draw a uniform time window centred around each respiration peak, with inspiration and expiration left and right of the peak, respectively (Fig. 1). EMG and MEG data were then segmented into ‘respiration-locked trials’ using the fixed time window. This data-driven approach allowed us to average across variations between respiration cycles and experimental runs while preserving the full dimensionality of individual respiration data.

**Fig. 1.**
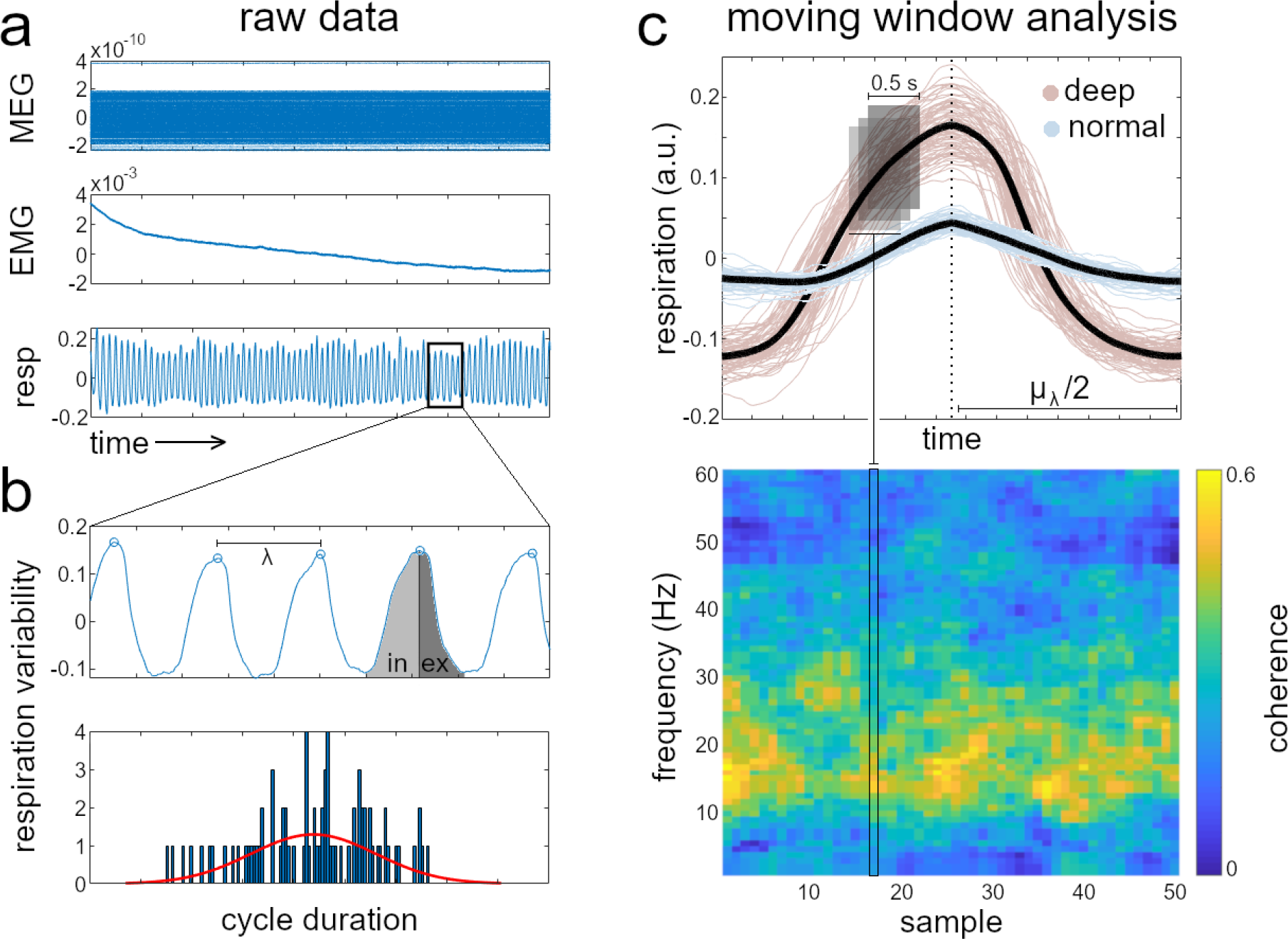
Data preprocessing and moving window analysis. **a** Segmentation of raw EMG and MEG data phase-locked to respiration. **b** Following identification of respiration peaks, the distribution of full-cycle durations (bottom) was used to generate a participant’s average respiration cycle for each respiratory condition and experimental run (in: inspiration; ex: expiration). **c** Moving window time-frequency analysis yielded CMC phase-locked to the respiration cycle. Top panel shows one participant’s average respiration cycles (black) plotted over single respiration cycles from the same run. Phase-locked segmentation nicely illustrates the increased tidal volume during deep (red) vs normal breathing (blue). While the duration of averaged respiration cycles varied between runs and participants, the selection of 50 equidistant samples across the individual respiration cycle allowed us to standardise phase information for further analyses across participants. Thus, resulting time-frequency representations of cortico-muscular coherence (bottom) were uniform in dimension and provided phase information of the entire respiration cycle of each trial.

For the calculation of time-frequency representations, we concatenated both runs of the same respiratory condition for each participant. An equal number of trials was used across conditions. Furthermore, variation in the duration of averaged respiration cycles was accounted for by sampling 50 equidistant time points over the course of each trial. A fixed time window with a duration of 0.5 s was centred on these time points in a moving window time-frequency analysis, precisely mapping spectral information across the entire respiration cycle (Fig. 1C). In this way, spectral power and cross-spectral density were calculated from 1 - 60 Hz (± 4 Hz smoothing) using a discrete prolate spheroidal sequences (DPSS) multitaper. Coherence between the EMG signal and all MEG channels within the beta band (10 - 30 Hz) was finally calculated for both respiratory conditions and averaged across participants.

### Statistical analyses

As outlined above, we first determined the location of maximum cortico-muscular coherence in sensor and source space based on continuous EMG and MEG data (i.e., not phase-locked to respiration). Based on previous research localising CMC (Schnitzler et al., 2000), we hypothesised sensors over the contralateral (i.e., left) sensorimotor cortex to primarily detect coherence in the beta band. Localisation of oscillatory sources was conducted using dynamic imaging of coherent sources (DICS; Gross et al., 2001) in Fieldtrip.

Using the ROI from this analysis, we first assessed CMC as a function of tidal volume. In other words, we aimed to illustrate how long-range, cortico-muscular communication within a well-established location depends on normal vs voluntary deep breathing. Therefore, coherence of EMG and MEG signals over left sensorimotor areas was contrasted with coherence over the right-hemispheric homologue by using a dependent-samples T-test to identify the frequencies for which left sensorimotor cortex showed significant CMC. Non-parametric cluster permutation tests (with 5000 randomisation iterations, significance level α = .05) were used to correct for multiple comparisons. In addition to independent testing of deep and normal breathing (vs zero, respectively), the direct contrast of deep vs normal breathing was to reveal the specific frequency band for which CMC was significantly enhanced (or reduced) over the entire respiration cycle. As stated in the introduction, we hypothesised tidal volume to modulate coherence within the beta band (10 - 30 Hz).

Our second hypothesis concerned modulatory effects of respiration phase on CMC within the beta band. Specifically, we employed circular-linear correlation between cortico-muscular coherence and respiration phase (using circ_corrcl.m in the CircStat toolbox for Matlab; Berens 2009) for all MEG sensors to detect significant cyclic modulation of CMC by respiration phase. Correlations were computed for a *single cycle* as well as *double cycle* model (assuming one and two full cycles over the time frame of 50 samples, respectively). This way, the single cycle model closely matched the respiration cycle whereas the double cycle model represented its first harmonic. We performed statistics across participants using dependent-samples T-test of fisher z-transformed correlations while correcting for multiple comparisons with cluster permutation tests (5000 randomisation iterations, α = .05).

## Results

### CMC in continuous data

Validating previous localisations of CMC in sensorimotor cortex (Brown, 2000; Gross et al., 2000; Kilner et al., 2000; Bourguignon et al., 2017), we first calculated coherence within continuous EMG and MEG data to extract a sensor space ROI for all further analyses. As hypothesised, the topographic distribution of CMC in the beta band (averaged from 10 - 30 Hz) was focussed over contralateral (i.e., left) medio-parietal sensors (Fig. 2a). Correspondingly, using the DICS beamformer algorithm for source reconstruction, we were able to localise the underlying neuronal sources within left sensorimotor cortex (Fig. 2b).

**Fig. 2.**
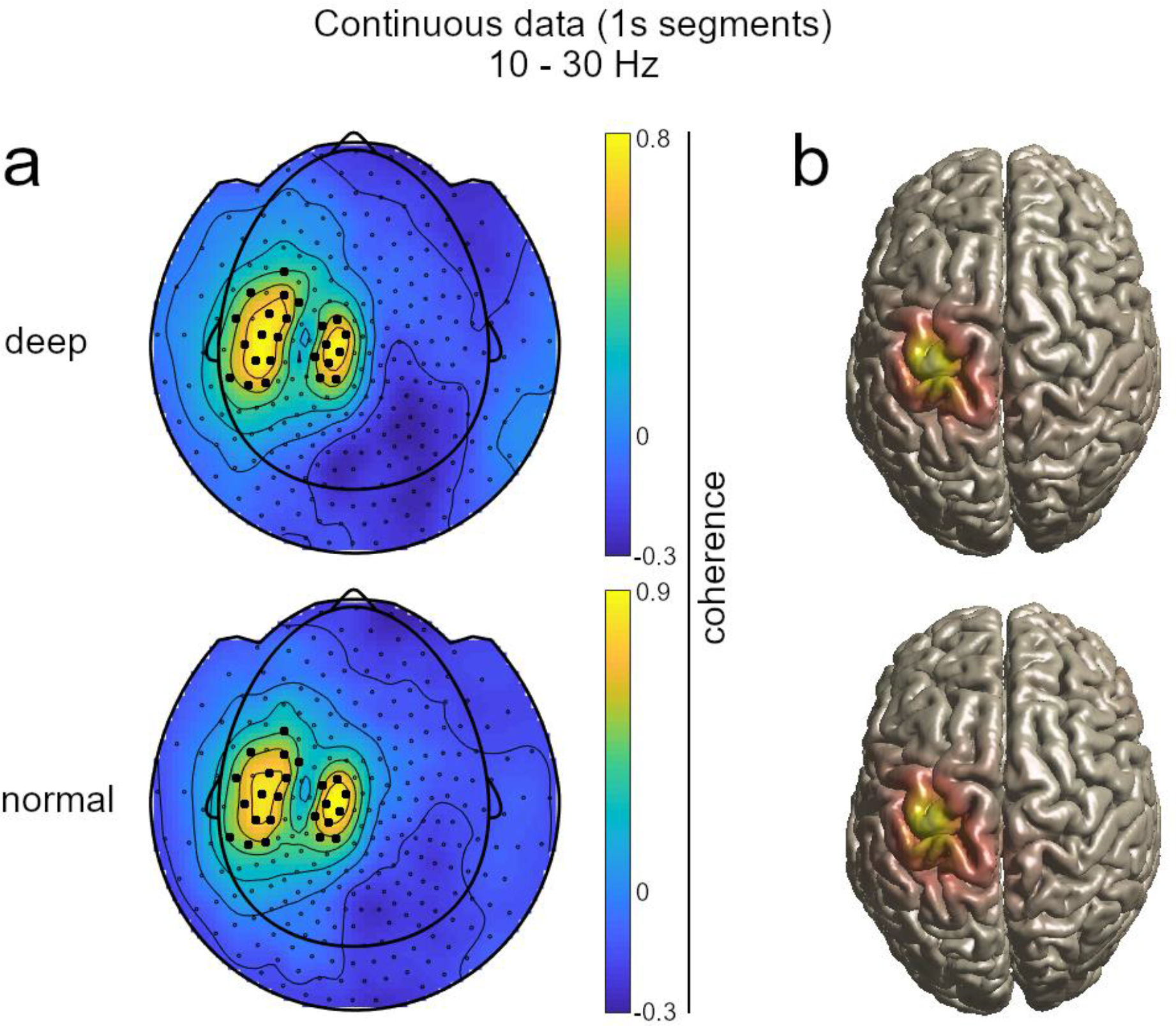
Group-level average of CMC in the beta range (10 - 30 Hz), calculated on 1 s segments of continuous data for deep (top row) and normal (bottom row) breathing. **a** Topographies were used to extract a sensor space ROI for all further analyses. Note that negative coherence values are due to spatial standardisation of individual coherence maps before averaging. **b** DICS localised the source of beta-range coherence in contralateral sensorimotor cortex.

### Coherence as a function of tidal volume

Looking at respiration-locked synchrony of EMG and MEG signals, our hypotheses regarding tidal volume effects were twofold: First, we assumed significant coherence of muscle and contralateral sensorimotor cortex activity in the beta range for both deep and normal breathing. Supporting this hypothesis, cluster permutation tests revealed significant CMC within the ROI for both respiratory conditions: For deep breathing, oscillatory neural activity between 10 - 33 Hz was found to be significantly synchronised with the EMG signal. For normal breathing, significant frequencies ranged from 6 - 34 Hz (Fig. 3).

**Fig. 3.**
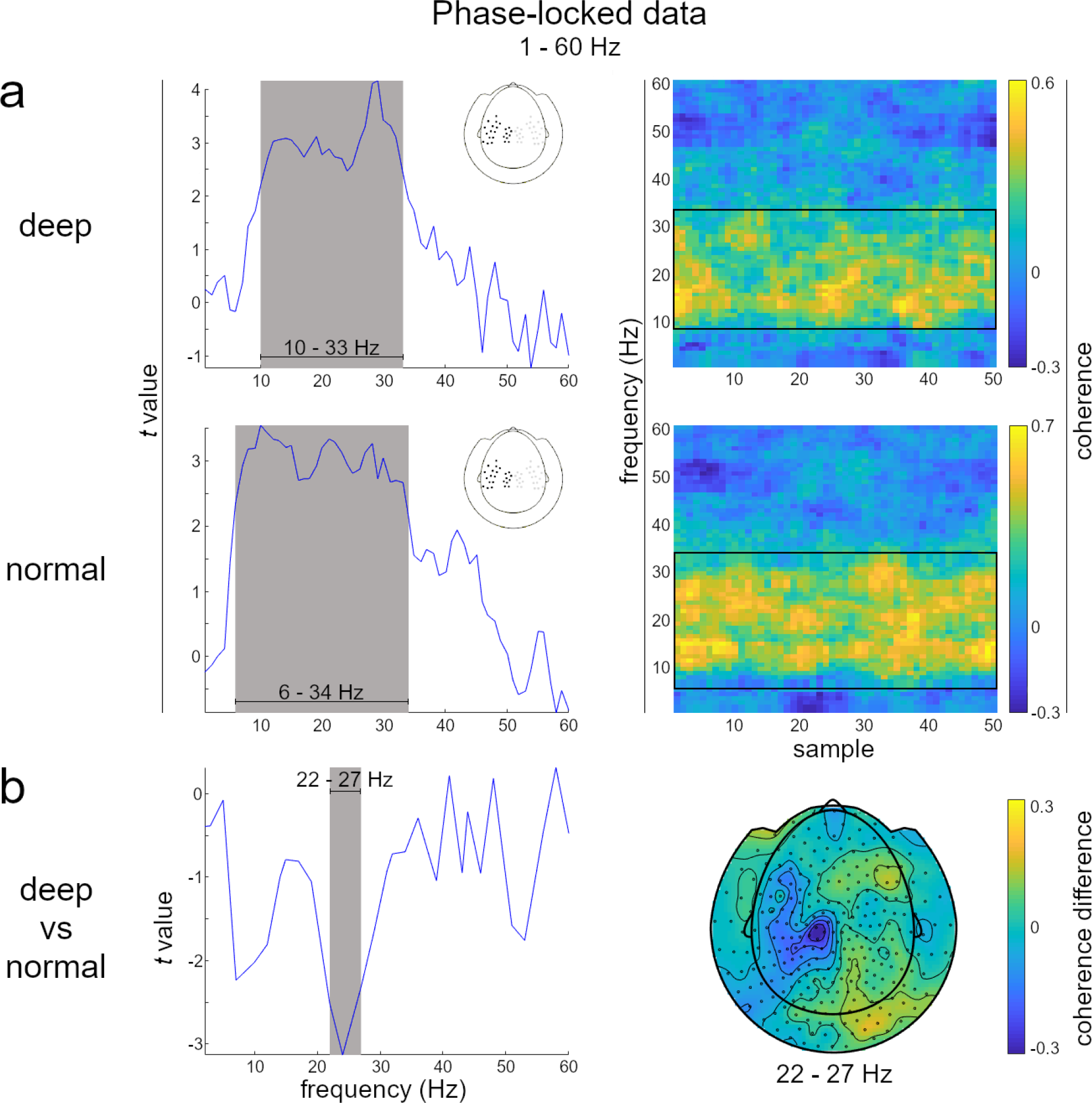
Coherence as a function of tidal volume. **a** Significant CMC was found for both deep and normal breathing within the ROI. Single conditions were tested against the contralateral homologue, respectively. Note that negative coherence values are due to spatial standardisation of individual coherence maps before averaging. **b** The contrast of deep vs normal breathing revealed significantly higher coherence in the normal breathing condition over the course of the entire respiration cycle.

Second, we hypothesised tidal volume to modulate CMC within the beta range. Supporting this hypothesis, a whole-head direct comparison of coherence (1 - 60 Hz) during deep vs normal breathing revealed a significant reduction of EMG-MEG synchrony at 22 - 27 Hz during voluntary deep breathing. This coherence reduction was localised predominantly at MEG sensors over the contralateral sensorimotor cortex (see Fig. 3).

### Coherence as a function of respiration phase

Adding to the global finding of overall changes in CMC (i.e., averaged across an entire respiration cycle) depending on tidal volume, our final hypothesis concerned coherence as a function of respiration phase. Specifically, we aimed to assess and localise beta-band CMC that was cyclically modulated over the course of a full respiration cycle. Since different complexities of such modulatory effects are conceivable, we tested whether beta coherence was significantly correlated with two separate cyclic signals: The single cycle model closely matched the course of the respiration signal, assuming one full cycle over the time frame of 50 samples that spans the respiration cycle (Fig. 4a). In contrast, the double cycle model assumed two full cycles over the same time frame, thus testing for cyclic modulation at a higher complexity. Critically, circular correlation computations for both models combined independently calculated orthogonal sine and cosine functions (Berens, 2009). This way, correlation of beta coherence and the respective cyclic signal was calculated at the optimal phase shift, effectively maximising the correlation coefficient.

**Fig. 4.**
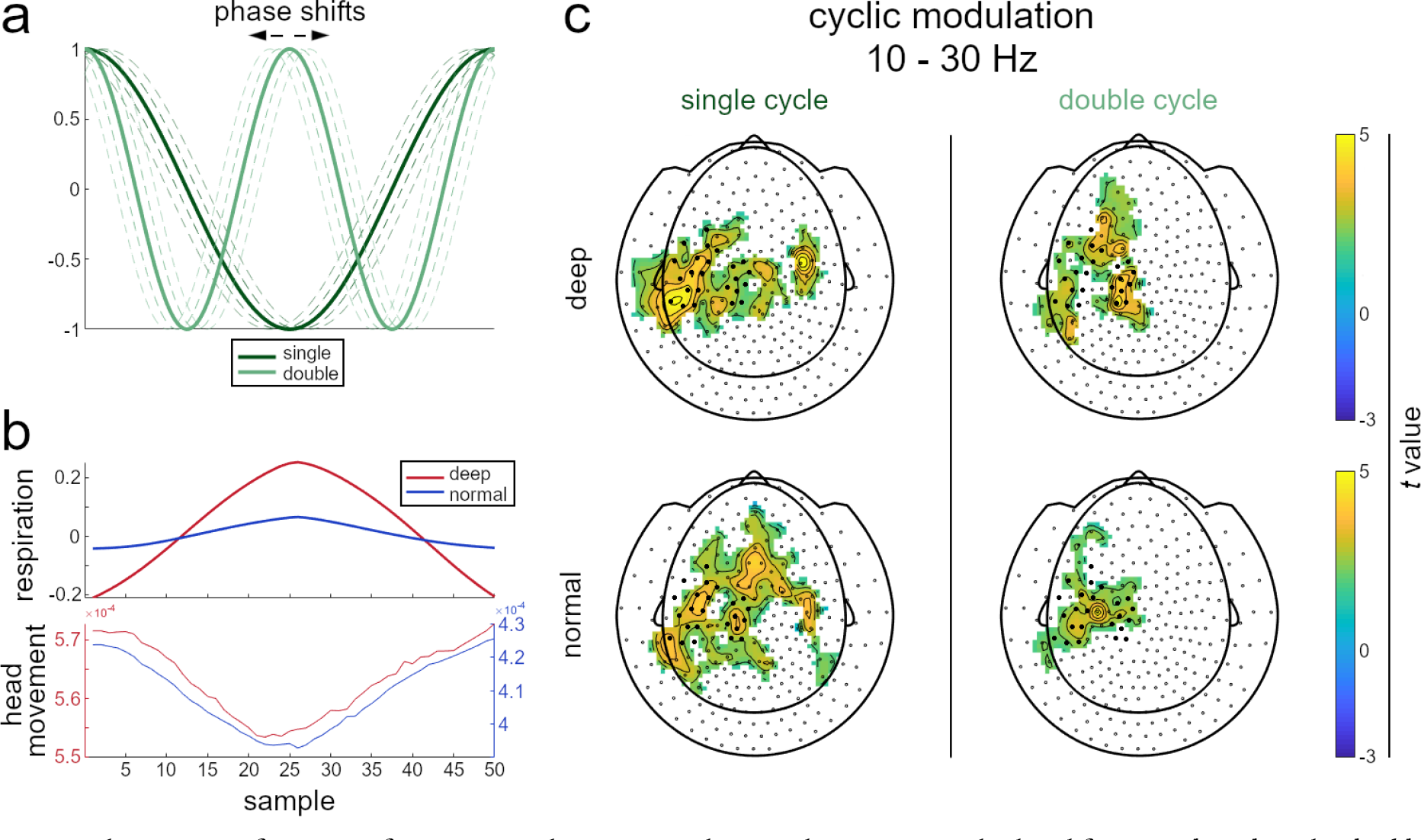
Coherence as a function of respiration phase. **a** Circular correlations were calculated for a *single cycle* and a *double cycle* model. Since these computations maximised correlations by means of orthogonal sine and cosine parts, they effectively return a correlation coefficient at the optimal phase shift (symbolised by dashed lines). **b** Group-level average respiration cycles for both respiratory conditions (top) as well as respiration-locked head movement (i.e., translation of the head centre relative to its starting position, bottom). The graph shows vector norms of translation in x, y, and z plane for deep and normal breathing. Note the difference in scaling due to expectedly increased head movement during deep (vs normal) breathing. **c** Topography of sensors where CMC (averaged from 10 - 30 Hz) was significantly cyclically modulated by respiration phase, according to the single cycle (left) and double cycle model (right). Circular correlation computations and cluster permutation tests were conducted on all sensors; ROI sensor positions are marked for illustrative purposes.

Substantiating our final hypothesis, cluster permutation testing revealed significant modulatory effects of respiration phase on EMG-MEG coherence (independent-samples T-test of fisher z-transformed correlations averaged from 10 - 30 Hz against 0). For both the single and the double cycle model, phase-locked modulation comprised - but was not restricted to - medio-parietal sensors contained in the ROI of our previous analyses, with a more extensive topography particularly found for the single cycle model (Fig. 4c). Taken together, these findings illustrate that i) respiration-locked modulation occurs at varying complexities which ii) each feature differential underlying topographies and iii) may, to a certain extent, be independent of the primary coherence amplitude per se.

### Coherence modulation and head movement

As expected we observed head movements phase-locked to the respiration cycle relative to the MEG sensors during measurements (measured as euclidean distance, see Fig. 4b). Moreover, increased thoracic circumference during deep breathing inevitably went along with increased breathing-related head movements. Therefore, when interpreting respiration-related modulation of corticomuscular coherence, it is instructive to consider the potential influence of head movement. In principle, breathing-related head movements can lead to rhythmic changes in the distance between MEG sensors and the participants head and thereby cause corresponding changes in the amplitude and signal-to-noise ratio of brain signals.

For respiration-related head movements to be the source of the reported modulation effects, changes in corticomuscular coherence would have to be significantly correlated to the respiration signal. To test this assumption, we computed linear Pearson correlation between the respiration signal and cortico-muscular coherence and subjected the results to cluster-corrected group analysis as described before. However, no significant effects were observed in normal or deep breathing condition. It should be noted, that for this analysis we intentionally used linear pearson correlation and not circular correlation because movement-related MEG artifacts will, first, monotonously follow respiration and movement time series, and second, do so with with zero lag (Figure 4b). Instead breathing-related neurophysiological effects might occur at non-zero time lags and with a temporal evolution different to the respiration time series while still being locked to the respiratory cycle. An example is the transient increase in the amplitude of high-frequency oscillations in the hippocampus during the short inspiration phase (Zelano et al., 2016).

To further strengthen the evidence against a confounding influence of head movements, we tested whether the single cycle model showed a significantly higher correlation with beta coherence than did the respiration signal. Recall that the single cycle model closely followed the respiration course (see Fig. 4a) but incorporated independent calculations of orthogonal sine and cosine functions in an angular-linear correlation. Hence, any incremental goodness of fit compared to the ‘raw’ respiration signal would be explained by the combination of the two functions, providing a solid argument against the influence of head movement. Consequently, we calculated a Wilcoxon signed rank test on the *t*-maps yielded by cluster permutation testing the correlation of beta-coherence with the respiration signal and the single cycle model. The test revealed a significantly better fit of the single cycle model for both deep (*Z* = −10.44, *p* < .001) and normal breathing (*Z* = −4.83, *p* < .001), rendering it highly unlikely that respiration-induced modulations in beta coherence were caused by concurrent head movement. Still, a note of caution is warranted for MEG studies in general. Since some aspects of human behaviour are modulated by respiration (see Introduction) any MEG studies investigating brain-behaviour relationships will benefit from simultaneously measuring respiration and using this information in their analysis.

## Discussion

The present study aimed to shed light on the fundamental but oftentimes neglected bi-directional interaction of brain oscillations and respiration as well as potential mechanisms underlying respiration-brain coupling. Specifically, we measured corticomuscular coherence in the beta band as a proxy of efficient long-range neural communication to characterise two potential modulatory mechanisms: First, we demonstrated overall higher cortico-muscular coherence during natural breathing compared to voluntary deep breathing. Second, we were able to show dynamic, cyclic modulation of beta coherence as a function of respiration phase. These findings critically underscore the explanatory potential of respiration-brain coupling regarding the orchestration of brain states and body states to optimise motor performance, emotion, perception, and cognition.

CMC originates from oscillatory activity in primary motor cortex that modulates firing probabilities of spinal motor neurons (Schoffelen et al., 2005). It leads to efficient cortico-peripheral coordination through at least three mechanisms that have been demonstrated in an in-vitro model of the rat respiratory system (Parkis et al., 2003). First, coherent oscillations reduce the variability of motor neuron spike trains. Second, they increase the gain (the number of action potentials that are elicited by a given input). Third, coherence makes the response of motor neurons more robust against changes in neurotransmitter levels. Therefore, CMC is a suitable marker for efficient cortico-peripheral communication.

With regard to the decrease of sensorimotor beta coherence during deep breathing, it is instructive to consider that increased tidal volume poses higher demands on respiratory muscles controlling, for example, thorax and diaphragm. The site of thoracic muscle representation has long been located in superior motor cortex, with the diaphragm representation immediately anterior thereto (Foerster, 1936). Indeed, previous studies have demonstrated enhanced motor activity during voluntary deep (vs automatic) breathing (McKay et al., 2003) as well as stronger respiration-locked oscillations in motor cortex during non-automatic (vs automatic) breathing (Herrero et al., 2018). Our finding of decreased sensorimotor beta coherence integrates these previous results in a comprehensive account of cortico-muscular (or cortico-spinal) communication: During voluntary, non-automatic respiration such as deep breathing, a distinct subset of M1 neurons targets spinal respiratory motoneurons as part of a cortico-motor loop that is not activated during automatic breathing (Evans et al., 1999; Murphy et al., 1990). Rather, automatic breathing is maintained by respiratory centres within the brain stem, primarily the preBötzinger complex (Del Negro et al., 2018), driving respiratory muscle activity (Tenney & Ou, 1977; Butler, 2007; Guz, 1997). In volitional control of respiration, however, bi-directional cortico-muscular signalling via fast-conducting corticospinal pathways is reflected in an additional ‘communication channel’ between motor cortex and respiratory motoneurons. Consequently, oscillatory activity within sensorimotor cortex now has to be synchronised separately with activity from both forearm and respiratory muscles, causing an overall decrease of beta coherence in motor areas during deep breathing. As for the functional implications of our two respiratory conditions, previous studies have employed motor evoked potentials (MEP) to demonstrate heightened corticospinal excitability in control of lower limb (Shirakawa et al., 2015) and finger muscles (Li & Rymer, 2011) during automatic (vs forced) breathing. Our results critically extend these results by showing that, on the cortical level, this enhancement is conceivably implemented by means of increased sensorimotor beta coherence during automatic breathing. Selective coherence has prominently been conceptualised as a general mechanism of neural interactions, regulating the efficiency of bi-directional communication through phase synchrony of input and output signals (Fries, 2005; Schoffelen et al., 2005).

In a similar vein, modulatory effects of respiration phase on beta coherence conceivably reflect dynamic variation in computational efficiency. A substantial body of literature has shown breathing rhythms to modulate oscillatory activity throughout the brain, leading to the suggestion of respiration as a common ‘clock’ in the organisation of neural excitability in the brain (Zelano et al., 2016; Kay 2005; Ito et al., 2014; Yanovsky et al., 2014). Together with consistent reports of respiratory adaptation in response to behavioural demands in animals (Welker, 1964; Kay & Freeman, 1998; Verhagen et al., 2007; Evans et al., 2009) and congruent evidence from human studies (Perl et al., 2019; Huijbers et al., 2014; Vlemincx et al., 2011), these findings strongly suggest an impact of respiration phase on sampling behaviour. While the continuous coupling of respiration and sampling behaviour is immediately obvious in animals’ sniffing, equivalent behavioural findings of spontaneous inhalation at task onset in humans have just gained attention fairly recently (Perl et al., 2019). Such spontaneous alignment of respiration phase to the sampling of sensory information in animals and humans uniformly points to an overarching optimisation mechanism: With cortical excitability varying over the respiration cycle, sampling information during phases of high excitability optimises efficient communication between brain areas and/or distal effector muscles. Conceptually, respiration has fittingly been cast as an example of *active sensing* (Corcoran et al., 2018) within the realm of predictive brain processing accounts (Mumford, 1992; Friston, 2005). This framework provides a comprehensive theoretical backdrop for respiration-brain coupling in that it underlines bi-directional signalling within cortico-cortical and cortico-muscular loops to maximise overall efficiency in neural processing. As outlined above, respiration conceivably actively entrains time windows of increased cortical excitability with the sampling of informative sensory stimuli. Mechanistically, the translation of slow neural rhythms into faster neural oscillations is widely regarded to be achieved through phase-amplitude coupling, as conclusively demonstrated in mice (Fries, 2005; Zhong et al., 2017; Ito et al., 2014). Respiration-induced rhythmic activity, transmitted through piriform cortex and subsequent cortico-limbic circuits (Fontanini & Bower, 2005; Fontanini & Bower, 2006; Litaudon et al., 2008), is thus a prime example of a slow oscillation whose phase rhythmically modulates the amplitude of faster oscillations. Hopefully, future studies will provide further insight into how respiratory rhythms actively modulate neural oscillations related to perception, cognitive processing, or even at rest.

Critically, previous findings of dynamic respiratory modulation, however illuminative, have so far been restricted to phase differences between inspiratory and expiratory breathing. In contrast, by means of circular correlation measures, our study is the first to provide novel insight into the complexities of respiration-related dynamics in neural communication. Specifically, we were able to show how CMC was cyclically modulated over the course of a respiration cycle as well as a signal modelling its first harmonic (represented by our single and double cycle models, respectively). As is always the case in model comparisons, a variety of other models can be conceived to characterise respiration-brain coupling. Consequently, while our findings show that binary accounts of respiratory cycles fall short of providing the full picture regarding coherence modulation, it remains an intriguing objective for future studies to characterise the shape of these modulatory processes in greater detail. It is our hope that the present study provides a starting point for the generation and stringent testing of hypothesis-based models in an effort to understand the interplay of respiration, brain oscillations, and behaviour.

## Acknowledgments

This study was supported by the Interdisciplinary Center for Clinical Research (IZKF) of the medical faculty of Münster (Gro3/001/19).

